# Circadian rhythm disruption is associated with skeletal muscle dysfunction within the blind Mexican Cavefish

**DOI:** 10.1101/2023.01.25.525368

**Authors:** Luke Olsen, Jaya Krishnan, Charles Banks, Huzaifa Hassan, Nicolas Rohner

**Affiliations:** Stowers Institute for Medical Research, Kansas City, MO 64110, USA; Department of Cell Biology & Physiology, University of Kansas Medical Center, Kansas City, KS 66160, USA

## Abstract

Circadian control of physiology and metabolism is pervasive throughout nature, with circadian disruption contributing to premature aging, neurodegenerative disease, and type 2 diabetes (Musiek et al. 2016; Panda, 2016). It has become increasingly clear that peripheral tissues, such as skeletal muscle, possess cell-autonomous clocks crucial for metabolic homeostasis (Gabriel et al. 2021). In fact, disruption of the skeletal muscle circadian rhythm results in insulin resistance, sarcomere disorganization, and muscle weakness in both vertebrates and non-vertebrates – indicating that maintenance of a functional muscle circadian rhythm provides an adaptive advantage. We and others have found that cavefish possess a disrupted central circadian rhythm and, interestingly, a skeletal muscle phenotype strikingly similar to circadian knock-out mutants; namely, muscle loss, muscle weakness, and insulin resistance (Olsen et al. 2022; Riddle et al. 2018; Mack et al. 2021). However, whether the cavefish muscle phenotype results from muscle-specific circadian disruption remains untested. To this point, we investigated genome-wide, circadian-regulated gene expression within the skeletal muscle of the *Astyanax mexicanus* – comprised of the river-dwelling surface fish and troglobitic cavefish – providing novel insights into the evolutionary consequence of circadian disruption on skeletal muscle physiology.

## Results/Discussion

We dissected skeletal muscle from adult surface fish and cavefish every 4 hours spanning a single daily cycle (24 hours) and sequenced the transcriptome via bulk RNA-sequencing (Fig. 1A). We identified 14,606 genes expressed in surface fish muscle and 14,723 genes expressed in cavefish muscle. To identify genes under circadian control, we utilized maSigPro and JTK_Cycle programs, two efficient algorithms for detecting rhythmic components in time-course experiments (Nueda et al. 2014; Hughes et al. 2010). Across both programs, we identified 440 and 370 circadian-regulated genes within the skeletal muscle of surface fish and cavefish, respectively – a 16% decrease in rhythmicity within cavefish muscle (Data S1). Confirming the accuracy of our pipeline, gene-ontology (GO) enrichment analysis revealed “circadian rhythm” and “rhythmic process” as the most enriched pathways within both surface fish and cavefish datasets (Fig. S1A and S1B), underscoring a cavefish muscle circadian rhythm, at least partially, remains intact. We identified multiple canonical clock genes within both surface fish and cavefish, such as *ciarta* and *per1a*. However, of these shared clock genes, almost all had changes in either amplitude or time of peak expression within cavefish relative to surface fish (Fig. 1B). Notably, the core clock gene *bmal1*, a crucial regulator of muscle composition and metabolism (Andrews et al. 2010), lacked rhythmicity within cavefish skeletal muscle, findings confirmed via qPCR (Fig. 1B and S1C).

**Figure 1.**
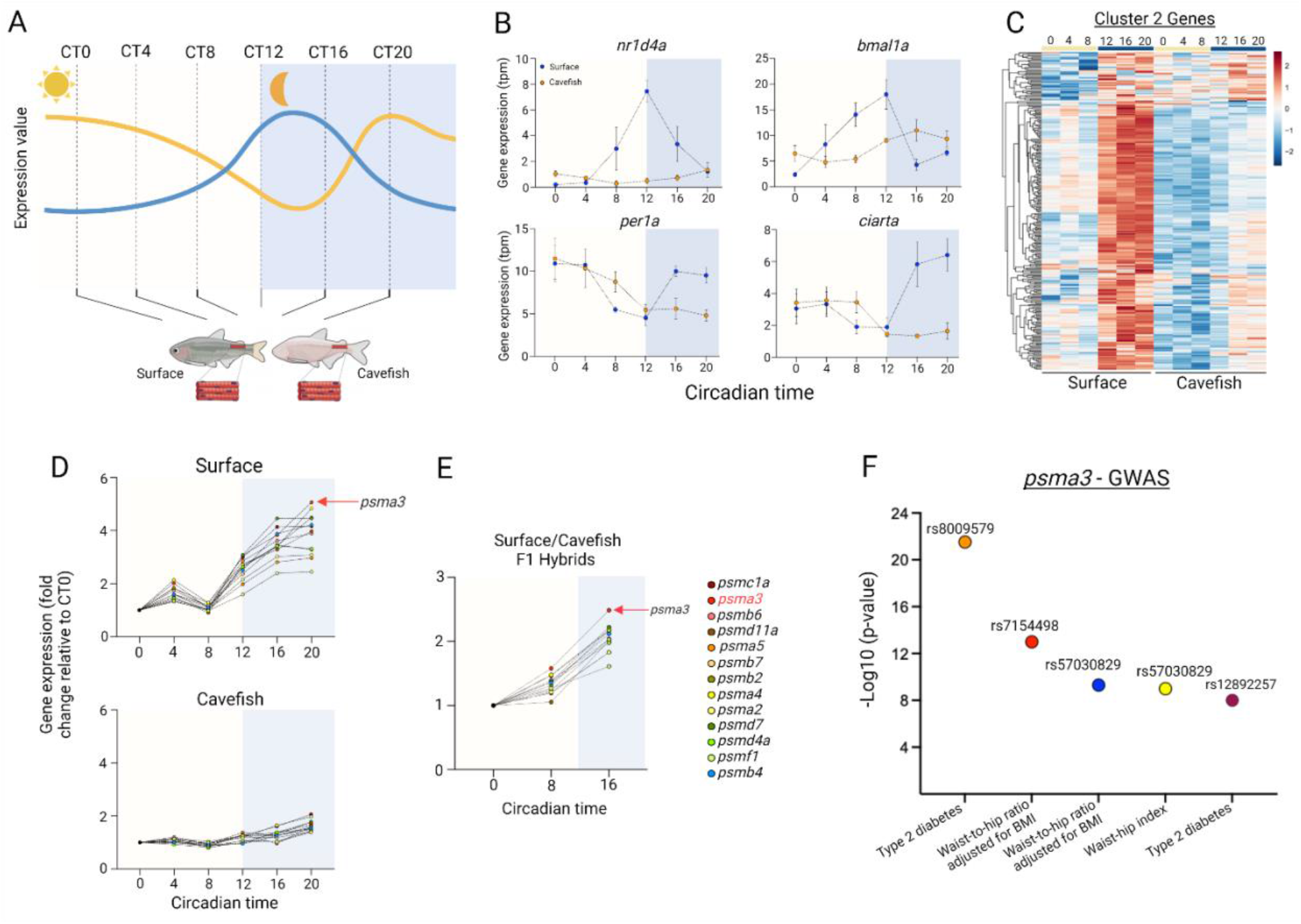
*Astyanax mexicanus* muscle circadian rhythm. (A) Schematic of tissue collection from the cavefish and surface fish. Timepoints are presented as circadian time (CT) with CT0 being 0600 and CT20 being 0200. For all analysis, a sample size of 3 for each morph and each timepoint were used. The orange and blue line are to represent the rhythmic (blue) and disrupted (orange) rhythmicity of a given gene. (B) Gene expression of various canonical clock genes between the surface fish (blue circles) and cavefish (orange circles). Data is presented as ±SEM. (C) Heatmap of all genes identified within the surface fish cluster 2 gene-set and the respective cavefish genes. (D) Circadian expression of proteasomal genes identified within the surface fish cluster 2 gene-set for (D) surface fish and cavefish and (E) cavefish/surface F1 hybrids. The red arrow places emphasis on *psma3*. (F) Single nucleotide polymorphisms (SNPs) of the *psma3* gene (including the psma3 antisense RNA 1) associated with an increase in the prevalence of obesity-related traits and Type 2 diabetes in humans (Data S5 and additional sourcing from https://www.ebi.ac.uk/gwas/genes/PSMA3). The variant/risk alleles are placed above each data point. All statistical analysis can be found within the supplementary material.

Having validated disruption of core clock genes within cavefish muscle, we sought to determine the effected muscle gene families – termed clock-controlled genes (CCGs). To do this, we performed k-means clustering of the 440 circadian regulated genes within surface fish muscle – here considered canonical *A. mexicanus* CCGs. We identified 4 distinct clusters peaking either during early (Circadian Time (CT) 0-4 – cluster 1 and 4), mid (CT 8 – cluster 3), or late (CT 16-20 – cluster 2) hours (Fig. S1D). Of the 4 clusters, ~60% of the genes fell into cluster 2, with GO-enrichment analysis revealing pathways enriched for muscle protein turnover, most notably the “proteasome complex” (Fig. S1E). Strikingly, ~93% of the genes within cluster 2 lacked rhythmicity within cavefish muscle – findings confirmed in a separate RNA-sequencing experiment (Fig. 1C and Fig. S2). Importantly, contrasting the cluster 2 genes against a recent whole-body circadian transcriptome of *A. mexicanus* (Mack et al. 2021) confirmed the cluster 2 gene-set oscillate in a muscle-specific manner (only 4 shared genes between datasets) and, subsequently, that their loss within cavefish is a muscle-specific phenomenon (Data S2).

The most striking example of cave-specific disruption in CCGs was that of the ubiquitin-proteasomal system (UPS). For example, multiple UPS genes peaked at night exclusively within surface fish muscle, most notably genes of the 20S proteasome catalytic core, the regulatory 19S proteasome, and various ubiquitin ligases, suggesting enhanced nocturnal circadian-regulated protein catabolism (Fig. 1D). In fact, proteomic analysis of muscle collected at CT0 and CT16 revealed surface fish had >3-fold more downregulated proteins at CT16 relative to CT0 as compared to cavefish, supporting increased nocturnal protein catabolism within surface fish muscle (Data S3).

To gain insight into the heritability of the proteasomal rhythmic phenotype, we performed bulk RNA-sequencing of skeletal muscle from cavefish-surface F1 hybrid’s collected at CT0, CT8, and CT16. We observed an intermediate nocturnal increase in proteasomal gene expression in the F1 fish (Fig. 1E), suggesting that the proteasomal gene rhythmicity is an incomplete dominant, potentially multigenic trait.

Considering the extreme loss in cavefish rhythmicity of the cluster 2 gene-set, we sought to determine possible transcriptional regulators. We conducted Ingenuity Pathway Analysis (Qiagen) and found *nfe2l2* (*Nrf2*) and *nfe2l1* (*Nrf1*) as top upstream regulators (*p*=3.96E-06 and *p*=6.96E-04, respectively) and, intriguingly, found both *nfe2l2* and *nfe2l1* were under circadian control exclusively within surface fish muscle (albeit only within our JTK_Cycle dataset). The Ingenuity pathway analysis further revealed Bmal, a transcriptional regulator of *nfe2l2* and *nfe2l1* (Early et al. 2018; Rey et al. 2011), as the most significantly upregulated “causal network” (*p*=2.08E-06) of the cluster 2 genes, suggesting circadian rhythmicity of Bmal transcriptionally regulates *nfe2l2* and *nfe2l1* expression and, subsequently, proteolytic gene expression – all of which are absent within cavefish muscle (Fig. S1F, S1G, and Data S4). Notably, querying the *bmal* sequence from wild-caught cavefish and surface fish revealed a cave-specific, non-synonymous mutation within its basic helix-loop-helix domain (human homolog: D93G) – a mutation residing two residues away from the documented Bmal1 disrupting L95E mutation (Huang et al. 2012) – a potential candidate underlying disrupted cavefish *bmal1* activity and subsequent cluster 2 gene-set rhythmicity.

Circadian control of protein turnover via proteasome processes is crucial for muscle quality, having been identified in mammalian and non-mammalian vertebrates (McCarthy et al. 2007; Kelu et al. 2020), with proteasomal activity necessary for amino acid availability, protein synthesis, and muscle growth. In fact, amino acid levels are under circadian control, with their rhythmicity coinciding with proteasomal gene expression (Eckel-Mahan et al. 2013). Recent metabolomic analysis from our group underscores this relationship, finding reduced cavefish muscle free amino acid levels coupled with decreased muscle mass and contractility (Medley et al. 2022; Olsen et al. 2022). For example, the anabolic amino acid leucine is decreased ~2-fold within cavefish skeletal muscle, with its transporter (*slc7a8*) lacking rhythmicity exclusively within cavefish (Fig. S1H). Interestingly, *slc7a8* similarly loses rhythmicity within muscle of aged mice and following exposure of human muscle cells to palmitate – conditions resulting in muscle dysfunction.

In agreement, contrasting genome-wide association studies (GWAS) against the disrupted cluster 2 gene-set revealed that ~23% of the 261 genes are associated with type 2 diabetes, obesity, and triglyceride levels – traits contributing to muscle atrophy and weakness (Data S5). In fact, the most arrhythmic proteasomal gene within cavefish muscle relative to both surface fish and cavefish-surface F1 hybrid’s (*psma3)* is associated with five GWAS SNPs for increased hip-to-waist ratio and type 2 diabetes in human populations (Fig. 1F). These findings highlight the evolutionary link between nocturnal protein turnover and muscle homeostasis, and reveal many known and novel candidate genes associated with circadian disruption and muscle function.

## Acknowledgments

We would like to thank the cavefish facility at the Stowers Institute for cavefish husbandry support, specifically Molly Miller, Elizabeth Fritz, David Jewell, Andrew Ingalls, and Diana Baumann. We thank all core support provided by the Stowers Institute; specifically, Rhonda Egidy and Amanda Lawlor of Sequencing and Discovery Genomics. We thank Suzanne McGaugh for providing the wild *A. mexicanus* genome sequences, Chris Seidel for computational analysis of the wild caught *Astyanax mexicanus* samples, and Alexander Kenzior for the initial identification of the *bmal1* mutation. NR is supported by institutional funding, NIH Grant 1DP2AG071466-01, NIH Grant R01 GM127872, NSF IOS-1933428, and NSF EDGE award 1923372.

---

**Supplementary Figure 1.**
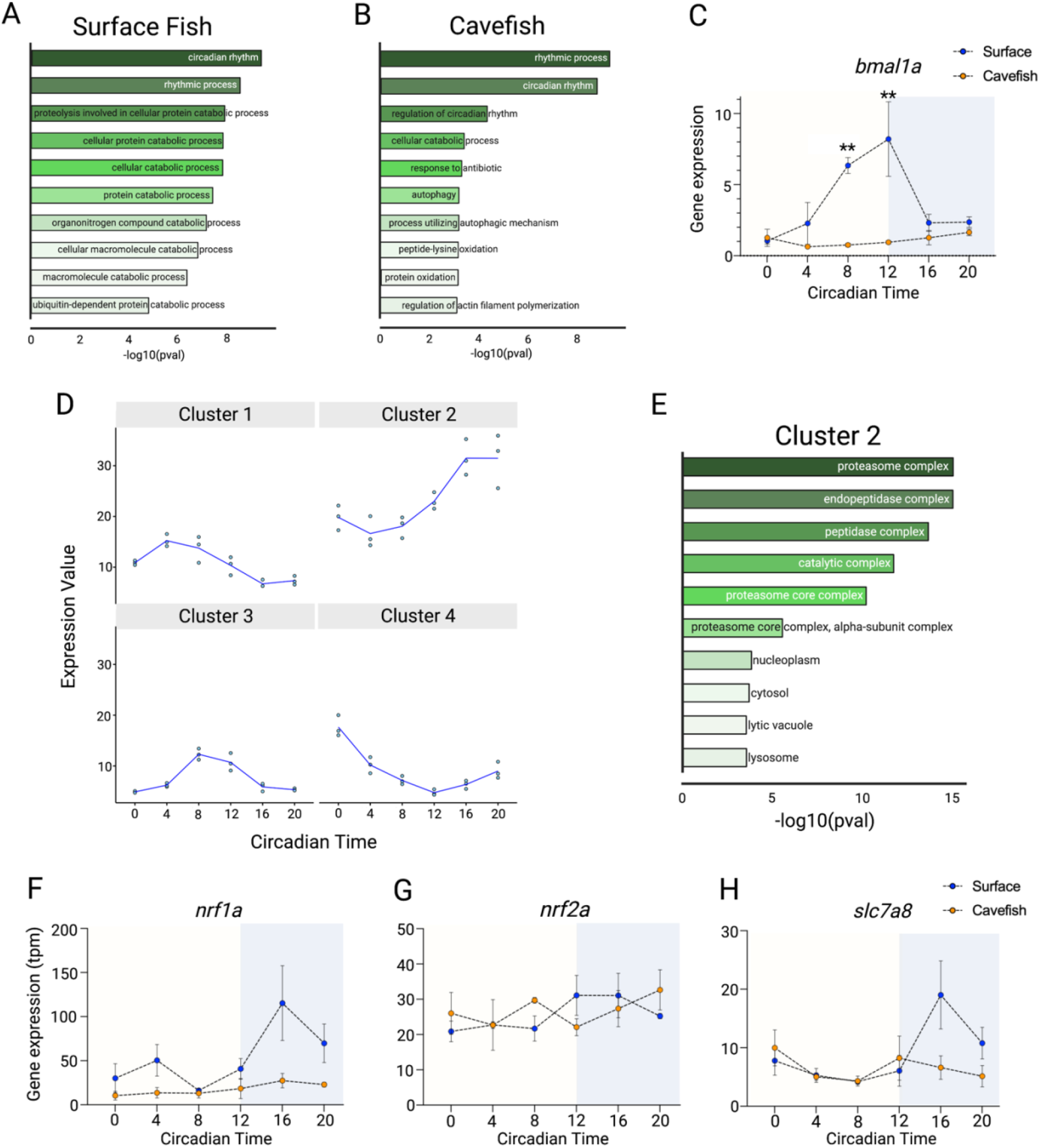
*Astyanax mexicanus* muscle circadian rhythm. Gene ontology enrichment analysis of circadian genes identified within JTK_Cycle and maSigPro within (A) surface fish and (B) cavefish. (C) Circadian rhythmicity of *bmal1a* via qPCR within cavefish and surface fish muscle (n=3 per morph/per timepoint). (D) The 4 clusters identified via k-means clustering analysis. Datapoints are the median gene expression. (E) GO-enrichment analysis of the 261 cluster 2 genes. Gene expression for (F) *nrf1a*, (G) *nrf2a*, and (H) *slc7a8* are expressed in transcripts per million (TPM). For figure S1C, a mixed-effects analysis with Benjamini & Hochberg correction was used. Data is presented as ±SEM, \*\**p*<0.01.

**Supplementary Figure 2.**
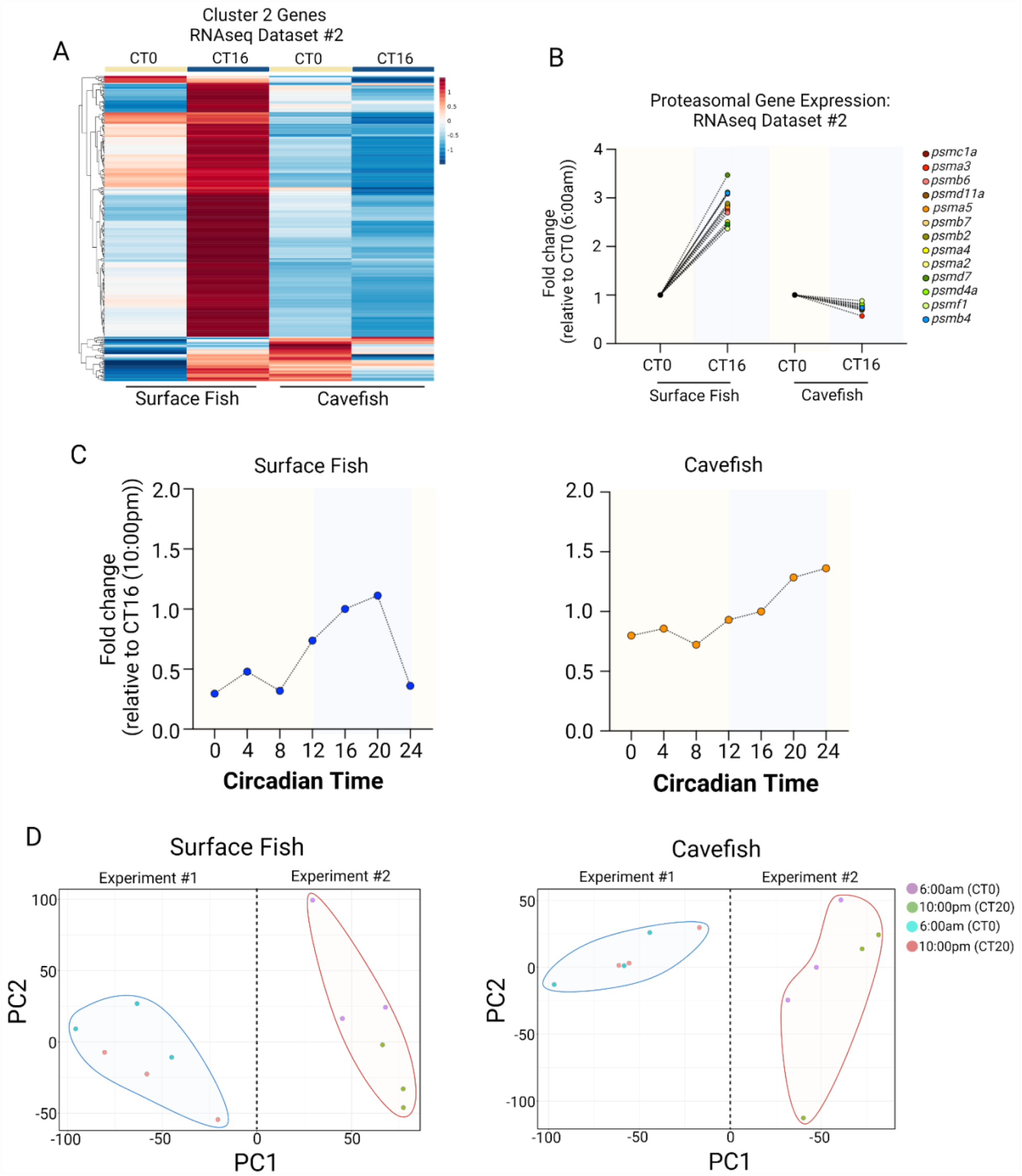
Validation of nocturnal rhythmicity within a separate cohort. (A) Heatmap of the cluster 2 genes identified within the initial experiment (Figure 1C). The presented data confirm the lack of nocturnal increase of these genes within cavefish skeletal muscle. (B) Fold-change of the proteasomal genes described in Fig. 1D. The CT16 timepoint is taken relative to the CT0 timepoint. (C) Additional 0600 timepoint (CT24) to confirm a rhythmic (24-hour) nature of the proteasome genes shown in Fig. 1D. To correct for the batch effect between RNA-sequencing experiments, both experiment 1 datapoints and experiment 2 datapoints were taken relative to the 2200 (CT16) samples within their respective runs. Shown is the mean of all proteasomal genes at each timepoint. (D) Principal component analysis of RNA-sequencing experiment 1 and experiment 2 including the 0600 (CT0) and 2200 (CT16) samples within surface fish and cavefish. These data are meant to highlight the batch effect between the respective RNA-sequencing runs, as can be seen by the individual experiments clustering together independent of time (CT0 vs CT16). These samples are further described in the Methods.

