## Supplemental Methods for "Circadian rhythm disruption is associated with skeletal muscle dysfunction within the blind Mexican Cavefish"

#### Laboratory fish husbandry

*Astyanax* husbandry and care was conducted identical to Olsen et al. (2022) and was approved by the Institutional Animal Care and Use Committee (IACUC) of the Stowers Institute for Medical Research on protocol 2021-122. All methods described here are approved on protocol 2021-129. Housing conditions meet federal regulations and are accredited by AAALAC International.

#### Lab fish muscle collection and RNA-sequencing

All fish used were adult and reared at similar tank densities. Laboratory-reared surface fish originating from the Rio Choy and cavefish originating from the Pachón cave were used throughout all studies. To confirm circadian rhythmicity of gene expression, fish were placed in complete darkness for 24 hours prior to muscle collection and were subsequently sampled every 4 hours within complete darkness, beginning at 0600 (CT0) and concluding at 0200 (CT20). Pachón/surface F1 hybrids were housed under identical conditions, however, the skeletal muscle was collected at 0600 (CT0), 1400 (CT08), and 2200 (CT16) under free-running (dark/dark) conditions. Similarly, an independent cohort of surface fish and cavefish were housed under identical conditions and their tissue was collected at 0600 (CT0) and 2200 (CT16) under free-running (dark/dark) conditions. These additional surface fish and cavefish samples were used to validate the primary findings of our initial experiment (described in figure 1). The data for the second, validatory experiment can be found in supplemental figure 2. For all fish, food was withheld on the day of muscle collection as to exclude all external stimuli. Fish were euthanized in 500mg/L of ms222 in complete darkness. Skeletal muscle was extracted immediately posterior to the dorsal fin and snap frozen in liquid nitrogen, all taking place in complete darkness. ~100mg of frozen tissue was homogenized in 1mL Trizol (Ambion) with triple pure M-Bio grade high impact zirconium beads in a Beadbug 6 microtube bead beater. RNA was extracted using standard phenol/chloroform extraction. The RNA pellet was cleaned with the RNeasy Mini Kit (Qiagen #74104) with on-column DNAse digestion (Qiagen #79256). Libraries were prepared according to the manufacturer's instructions using the TruSeq Stranded mRNA Prep Kit (Illumina #20020594). The resulting libraries were quantified using a Bioanalyzer (Agilent Technologies) and Qubit fluorometer (Life Technologies). Libraries were normalized, pooled, multiplexed, and sequenced on an Illumina NextSeq 2000 instrument as v2 Chemistry High Output 100bp single read runs. Following sequencing, raw reads were demultiplexed into Fastq format allowing up to one mismatch using Illumina bcl2fastq2 v2.20. Reads were aligned to UCSC genome astMex_2.0 with STAR aligner (version 2.7.3a) using Ensembl 102 gene models. TPM values were generated using RSEM (version v1.3.0). Pairwise differential expression analysis was performed using Bioconductor package edgeR (3.36.0 with R 4.1.2). Only protein coding genes and long non-coding RNAs (lncRNAs) were considered from the Ens_102 annotation. Only genes with counts per million expression of ≥2 in at least 2 samples were kept for further analysis. Statistical significance was determined by fold change cutoff of 2 and false discovery rate (FDR) cutoff of 0.05 with the p.adjust function in R with the default choice Benjamini & Hochberg correction. Gene Ontology (GO Term) enrichment was completed using TERMS2GO, an in-house R Shiny app (versions R 4.1.0, shiny 1.7.1). Significant gene ontology terms were identified using clusterProfiler's enrichGO function (version 4.0.0) with Annotation Hub's species database (version 3.0.0). GO Terms with adjusted p-values less than 0.05 were considered significant. Figures were generated using ggplot2 (version 3.3.5) and plotly (4.10.0). Ingenuity Pathway Analysis was conducted with the use of QIAGEN IPA (QIAGEN Inc., https://digitalinsights.qiagen.com/IPA) (Krämer et al. 2014).

**Time series analysis**

Rhythmic genes were identified using the R packages 'JTK_cycle' (v 3.1) (Hughes et al. 2010) and maSigPro (v 1.66) (Nueda et al. 2014). Gene counts were normalized using trimmed mean of M values (TMM) method in edgeR (v 3.36) (Robinson et al. 2010). JTK_cycle was run on a 24-hour cycle with a spacing of 4 hours and 3 replicates per time point. Amplitude and Bonferroni-adjusted p-values were also computed using JTK_Cycle. Genes with adjusted p-value < 0.05 were retained as significant rhythmic genes. maSigPro was run using the Generalized linear model (GLM) setting and p-value's corrected with Benjamini & Hochberg correction. Genes at FDR 5% were retained as significant rhythmic genes. Significant genes identified within both cycling programs were grouped into k = 4 groups using k-means clustering method.

**Proteomics**

Skeletal muscle from adult surface fish and cavefish (Pachón) were collected under free-running (dark/dark) conditions as described above. Fish were collected at 0600 (CT0), 1400 (CT08), and 2200 (CT16). Skeletal muscle was immediately snap frozen in liquid nitrogen. ~30mg of skeletal muscle was homogenized in 800µl of RIPA Buffer (ThermoScientific) supplemented with protease inhibitor and homogenized with triple pure M-Bio grade high impact zirconium beads in a Beadbug 6 microtube bead beater. Samples were then placed at 4˚C for 1.5 hours and gently rocked. Samples were subsequently centrifuged at 16,900 x rpm at 4˚C for 20 minutes. The supernatant was collected, and protein was measured with the BCA Protein Assay Kit (ab102536). 100µg of protein was then precipitated using the Tricarboxylic Acid Precipitation and washed twice with ice cold acetone. Dried protein pellets were resuspended in buffer containing 100 mM Tris-HCl, pH 8.5 and 8M urea, incubated with 5 mM TCEP (tris(2-carboxyethyl)phosphine) at room temperature for 30 minutes, incubated with 10 mM CAM (2-chloroacetamide) at room temperature for 30 minutes, and digested with 0.1 μg endoproteinase Lys-C for 6 hours at 37°C. The concentration of urea was reduced to 2 M using 100 mM Tris-HCl, pH 8.5. $CaCl_2$ was added to 2 mM final concentration. Samples were incubated with 0.5 μg trypsin at 37°C overnight and reactions stopped with formic acid (5% final concentration). The resulting peptides were desalted using Pierce™ Peptide Desalting Spin Columns (#89852) according to the manufacturer's instructions, quantitated using the Pierce™ Quantitative Fluorometric Peptide Assay, and 5µg of peptide was resuspended in buffer A (5% acetonitrile, 0.1% formic acid) prior to mass spectrometry analysis. Multidimensional protein identification technology (MudPIT) was performed essentially as described previously (Swanson et al. 2009) using an Orbitrap Elite mass spectrometer. Resulting .raw files were converted to .ms2 files using the in-house software package RAWDistiller v. 1.0. The ProLuCID search algorithm was used to match spectra to a database containing 39383 *Astyanax mexicanus* protein sequences NCBI (NCBI 02/26/2021 release) and 426 contaminant protein sequences. Mass tolerances were set at 10 ppm for precursor ions and 500 ppm for fragment ions. A static modification of 57.02146 Da was added to cysteine residues to account for carboxamidomethylation; variable modifications of 114.04293 Da for lysine residues (ubiquitination) and 15.9949 Da for methionine residues (oxidation) were also searched. The result files from the ProLuCID search engine were processed with DTASelect (v 1.9) (Tabb et al. 2002). The in-house software package swallow (v 0.0.1), which works with DTASelect, was used to select Peptide Spectrum Matches so that the false discovery rates (FDRs) at the peptide and protein levels were less than 5%. Peptides and Proteins detected across all samples were compared using CONTRAST (Tabb et al. 2002). Proteins that were identified by the same set of peptides (including at least one peptide unique to such protein group to distinguish between isoforms) were grouped together, and one accession number was arbitrarily considered as representative of each protein group. Protein abundances in experimental and control samples were assessed by calculating dNSAF values (Zhang et al. 2010). The QPROT statistical framework was used to assess protein differences between sample groups (Choi et al. 2015).

**RT-qPCR**

RNA was used from the samples collected for RNA-sequencing as described above. 1-5ng of cDNA was used for qPCR utilizing SYBR green technology (Quantabio PerfeCTa SYBR Green FastMix Low Rox #66188573) with a QuantStudio 5 Real-Time PCR System. Specificity of each amplicon was confirmed via analysis of post-reaction dissociation curves, validating a single amplicon for each set of primers. Analysis was conducted using the Delta Delta $C_t$ method. All samples were run in triplicate and normalized to the housekeeping gene *rpl13a*.

Primer sequences used are as follows:

*Bmal1a:*

FW: 5'- ATGCCAAAACTGGTCTGCCT -3'

RV: 5'- TCCATCTTGACGGACGGTTG -3'

*rpl13a:*

FW 5'- GTTGGCATCAACGGATTTGG -3'

RV: 5'- CCAGGTCAATGAAGGGGTCA -3'

**Bmal1 mutation analysis**

We used bcftools to query the nucleotide sequence at chr24 position 26625383 in a VCF file kindly provided by Suzanne McGough, built from genomic reads of wild-caught Pachón and Tinaja cavefish and wild-caught surface fish from the Rio Choy and Rascón localities using GATK. We further confirmed the presence of the T-to-C mutation using samtools mpileup with BAM files from ATAC-Seq datasets performed on laboratory-reared Rio Choy surface fish and Pachón and Tinaja cavefish.

### Data Availability

The following secure token has been created to allow review or record GSE196531 while it remains in private status: ebgxgcaezpkhngb. All proteomic data has been uploaded to MassIVE repository with identifier MSV000091062 and can be accessed with password “circadian”. Original data underlying this manuscript may be accessed after publication from the Stowers Original Data Repository.
